# Ebola virus exploits host lncRNA *LINC01740* to enhance ATF3 and suppress antiviral immune responses

**DOI:** 10.64898/2026.07.06.731958

**Authors:** Olena Shtanko, Tanuj Gunturu, Anu Gopal, Marija Djurkovic-Lopez, Hoang Nguyen, Sahana Jayakumar, Ajai Lawrence D’Silva, Asha Thomas, Smita Kulkarni

## Abstract

Ebola virus (EBOV) infection causes severe hemorrhagic fever marked by dysregulated cytokine production, impaired antiviral defenses, and multi-organ failure. Macrophages are primary targets of EBOV, and viral replication profoundly alters macrophage transcriptional programs, driving hyperinflammation. Although long non-coding RNAs (lncRNAs) are increasingly recognized as regulators of immunity and viral pathogenesis, their roles in EBOV infection remain poorly understood. We performed comprehensive transcriptomic profiling of primary human monocyte-derived macrophages infected with the highly pathogenic EBOV Mayinga variant. Infection triggered extensive remodeling of both coding and non-coding transcriptomes, including hundreds of differentially expressed lncRNAs. Functional analysis of neighboring protein-coding genes of EBOV-induced lncRNAs (EVILs) revealed enrichment of pathways linked to cytokine signaling, transcriptional regulation, and cell signaling, all of which are central to Ebola virus disease (EVD) pathogenesis. Among the most strongly induced EVILs, LINC01740 and its neighboring protein-coding gene, Activating Transcription Factor 3 (ATF3), were significantly upregulated. Antisense oligonucleotide-mediated inhibition of LINC01740 reduced ATF3 mRNA and protein levels. CRISPR/Cas13d-mediated knockdown of ATF3 restored type I interferon (IFN-I) signaling and antiviral gene expression in EBOV-infected macrophages. Mechanistically, ATF3 functions as a negative regulator of IFN-I and type I interferon-stimulated gene expression, thereby suppressing antiviral immune responses in EBOV-infected macrophages. Together, these findings identify a previously unrecognized LINC01740-ATF3-IFN-I regulatory axis that EBOV exploits to promote immune suppression and viral replication.

## Introduction

Ebola virus (EBOV) causes a severe hemorrhagic disease with high case fatality rates and recurrent outbreaks in sub-Saharan Africa. Despite repeated epidemics, including the 2025 outbreak in the Democratic Republic of Congo, which had a case fatality exceeding 60%[1], FDA-approved countermeasures remain limited. Ebola virus disease (EVD) is clinically characterized by acute fever, hypotension, shock, and profound coagulation abnormalities. The extraordinary pathogenicity of EBOV stems from its ability to efficiently infect target cells while suppressing innate immune responses early in the viral replication cycle [2]. Experimental studies in nonhuman primates have demonstrated that macrophages and dendritic cells (DCs) are primary early cellular targets of infection [3]. EBOV replication in macrophages generates high titers of progeny virions that disseminate systemically, inducing cellular reprogramming that drives robust production of proinflammatory mediators. This dysregulated immune activation contributes to vascular leakage, coagulopathy, multi-organ failure, and death [4]. Uncontrolled EBOV replication drives macrophage dysfunction and excessive cytokine release, resulting in a mixed pro- and anti-inflammatory “cytokine storm.” Elevated levels of vasoactive peptides, tissue factor, and tumor necrosis factor promote hypotension and disseminated intravascular coagulation, both hallmarks of severe EVD [2, 4, 5]. Rather than conferring protection, EBOV-induced macrophage activation appears to amplify disease pathogenesis by facilitating viral spread and fueling injurious inflammation [3]. However, the mechanisms by which EBOV orchestrates this maladaptive host response remain incompletely understood.

Only about 2% of the human genome encodes proteins; the majority of the transcriptome consists of non-coding RNAs, including long non-coding RNAs (lncRNAs), which have emerged as critical regulators of gene expression. lncRNAs modulate diverse cellular processes, including antiviral defense, inflammatory signaling, and viral replication[6–16]. They regulate the transcription of protein-coding genes, influence protein localization and stability, and mediate interactions among proteins and nucleic acids. Host lncRNA expression is markedly altered in multiple viral infections, suggesting that these transcripts form extensive regulatory networks that expand the complexity of immune gene regulation and may shape disease outcomes [9, 17–25]. Recent single-cell analyses in EBOV-infected macaques identified numerous infection-responsive lncRNAs [26]. Many lncRNAs are co-expressed with immune-regulatory protein-coding genes or change upon viral entry in EBOV-infected macaques, suggesting a functional role in host–virus interactions [26]. Notably, investigation of human non-coding RNA functions in EBOV-infected cells has been limited to a tetracistronic transcription- and replication-competent virus-like particle (trVLP) system in 293T cells, leaving a critical gap in our understanding of the functional impact of lncRNA responses to wild-type EBOV in primary human immune cells [27].

To address this gap, we characterized changes in lncRNA and protein-coding mRNA expression in primary human monocyte-derived macrophages infected with the highly pathogenic wild-type EBOV variant Mayinga [28]. Our analyses revealed extensive remodeling of the macrophage lncRNA landscape during infection and identified a subset of EBOV-induced lncRNAs (EVILs) with previously uncharacterized roles in cellular processes and EBOV pathogenesis. Among these, we demonstrate that EBOV robustly induces Activating Transcription Factor 3 (ATF3) early after infection. Silencing ATF3 enhanced innate immune signaling and antiviral gene expression, indicating that EBOV exploits ATF3 to dampen macrophage antiviral responses and delay immune activation. We further show that LINC01740 is significantly upregulated in infected macrophages and augments ATF3 expression. Together, these findings uncover a previously unrecognized lncRNA-ATF3 regulatory axis that contributes to immune suppression and viral propagation during EBOV infection.

## Results

### EBOV infection of human primary macrophages causes significant global changes in mRNA and lncRNA expression

Given the vast potential for discovering novel mechanisms of immune response control by lncRNAs, we sought to determine the non-coding transcriptome of primary human macrophages infected with EBOV and to develop a model system for functional analyses of lncRNA regulatory processes. A pool of cells from nine donors, including both males and females, served as the source of monocyte-derived macrophages (MDMs). We infected the MDMs with the EBOV variant Mayinga at a multiplicity of infection (MOI) of 1 and analyzed the cellular transcriptome at 24 h and 48 h post-infection. We identified significant temporal changes in the expression of lncRNAs (Fig. 1a, b) and protein-coding genes (Fig. 1c, d) in EBOV-infected human macrophages. Differential expression analysis identified 576 lncRNAs that were significantly upregulated and 36 lncRNAs that were significantly downregulated in EBOV-infected macrophages compared with mock-infected controls at 24 h post-infection. Similarly, among protein-coding transcripts, 338 mRNAs were significantly upregulated, and 29 mRNAs were significantly downregulated in EBOV-infected vs. Mock-infected macrophages (adjusted p < 0.05, fold change ≥ 2; Fig. 1). We performed Ingenuity Pathway Analyses (IPA) on differentially expressed genes identified by transcriptomic profiling of EBOV-infected vs. Mock-infected macrophages at 24 h post-infection. IPA analyses showed robust activation of innate immune and inflammatory signaling programs in EBOV-infected macrophages (Supplementary Fig. S1).

**Figure 1.**
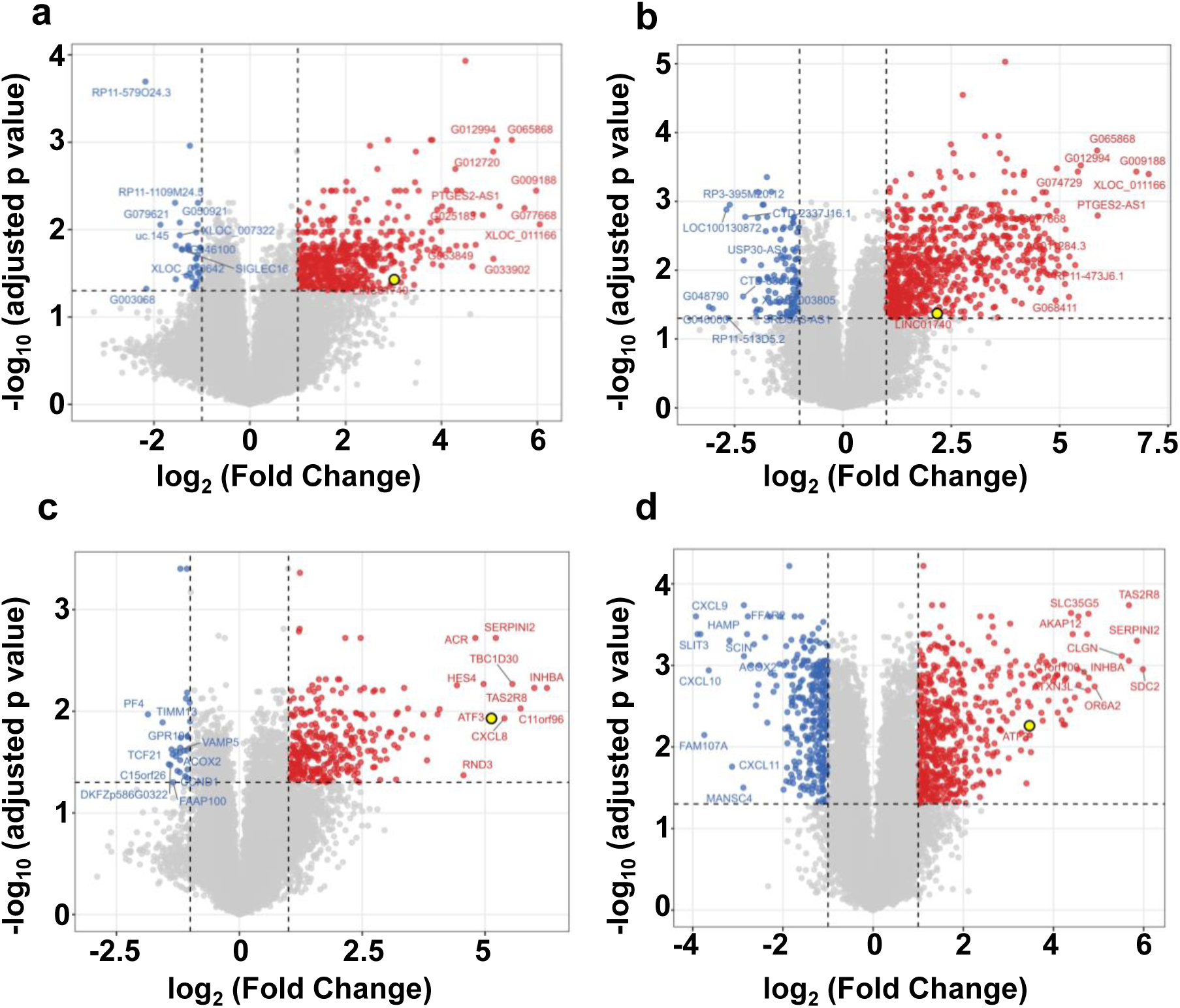
EBOV infection dynamically reshapes lncRNA and mRNA expression in primary human macrophages. Volcano plots show differential expression of lncRNAs **(a and b)** and protein-coding mRNAs **(c and d)** in EBOV-infected versus mock-infected primary human macrophages at 24 h **(a and c)** and 48 h **(b and d)** post-infection. The x-axis shows log₂ fold change (log_2_FC), and the y-axis shows −log_10_ adjusted p-value. Total RNA from three independent biological replicates per condition was analyzed by microarray. Genes with a Benjamini-Hochberg false discovery rate (FDR)-adjusted p-value of <0.05 and an absolute log_2_FC of ≥|1| (≥|2|-fold change) were considered differentially expressed. Significantly upregulated and downregulated genes are shown in red and blue, respectively.

LncRNAs regulate neighboring protein-coding genes through diverse mechanisms [29, 30]. Genome-wide transcriptomic analyses have revealed that many lncRNAs are co-expressed with neighboring protein-coding genes, consistent with potential cis-regulatory relationships, although such correlations do not necessarily imply functional interactions[31–34]. Because functional validation has been performed only for a few lncRNAs, several studies apply the ‘guilt-by-association’ principle to infer lncRNA functions on a genome-wide scale [35]. We adapted this approach to understand the distinct co-regulation of EVILs and protein-coding mRNAs. We characterized the genomic distribution of EVILs, identified the nearest protein-coding neighbor within 300 kb, and performed functional enrichment analysis of the nearest-neighbor protein-coding genes. Differentially expressed protein-coding mRNA neighbors of highly induced EVILs are involved in cytokine production and cytokine-mediated signaling, cell signaling pathways, cell proliferation, apoptosis, and most significantly, transcriptional regulation (Supplementary Fig. S2). Several of these proteins have been previously implicated in EVD pathology [5, 36, 37]; however, the roles of others, such as cell cycle regulation, apoptosis, and cell signaling pathways, are poorly characterized [36, 38–41]. LncRNAs are highly responsive to pathogenic stimuli [9, 17–19, 21–24, 42–44] and frequently act in *cis* to regulate neighboring protein-coding genes. Functional *cis*-acting lncRNAs are especially enriched around genes encoding transcription machinery [45]. Given the enrichment of EVILs near transcriptional regulators, we hypothesized that select EVILs are induced during EBOV infection and coordinate the expression of adjacent transcription factors.

### ATF3 and its neighboring lncRNA, LINC01740, are co-regulated in EBOV-infected MDM

Our data show that EBOV infection induces rapid global changes in the macrophage transcriptome (Fig. 1). Among these changes, a significant increase in ATF3 expression in EBOV-infected human primary macrophages is consistent with other reports [36, 46].

One of the EVILs, LINC01740, is co-regulated with its neighboring protein-coding gene, ATF3. LINC01740 is located upstream of ATF3 (Fig. 2a). We refer to LINC01740 as the ‘untranslated neighbor of ATF3’ (UNA) throughout the rest of the manuscript. UNA expression is significantly higher in EBOV-infected macrophages than in mock-infected cells (Fig. 2b). mRNA levels were measured by quantitative PCR (qPCR) using gene-specific primers (Supplementary Table S1). mRNA and protein levels of ATF3 are also markedly elevated in EBOV-infected macrophages compared with mock-infected cells (Fig. 2b, c). A published report found that both UNA and ATF3 mRNA expression levels are significantly higher in Makona variant-infected THP-1 macrophages than in uninfected cells, corroborating our findings (Supplementary Figure S3) [47].

**Figure 2.**
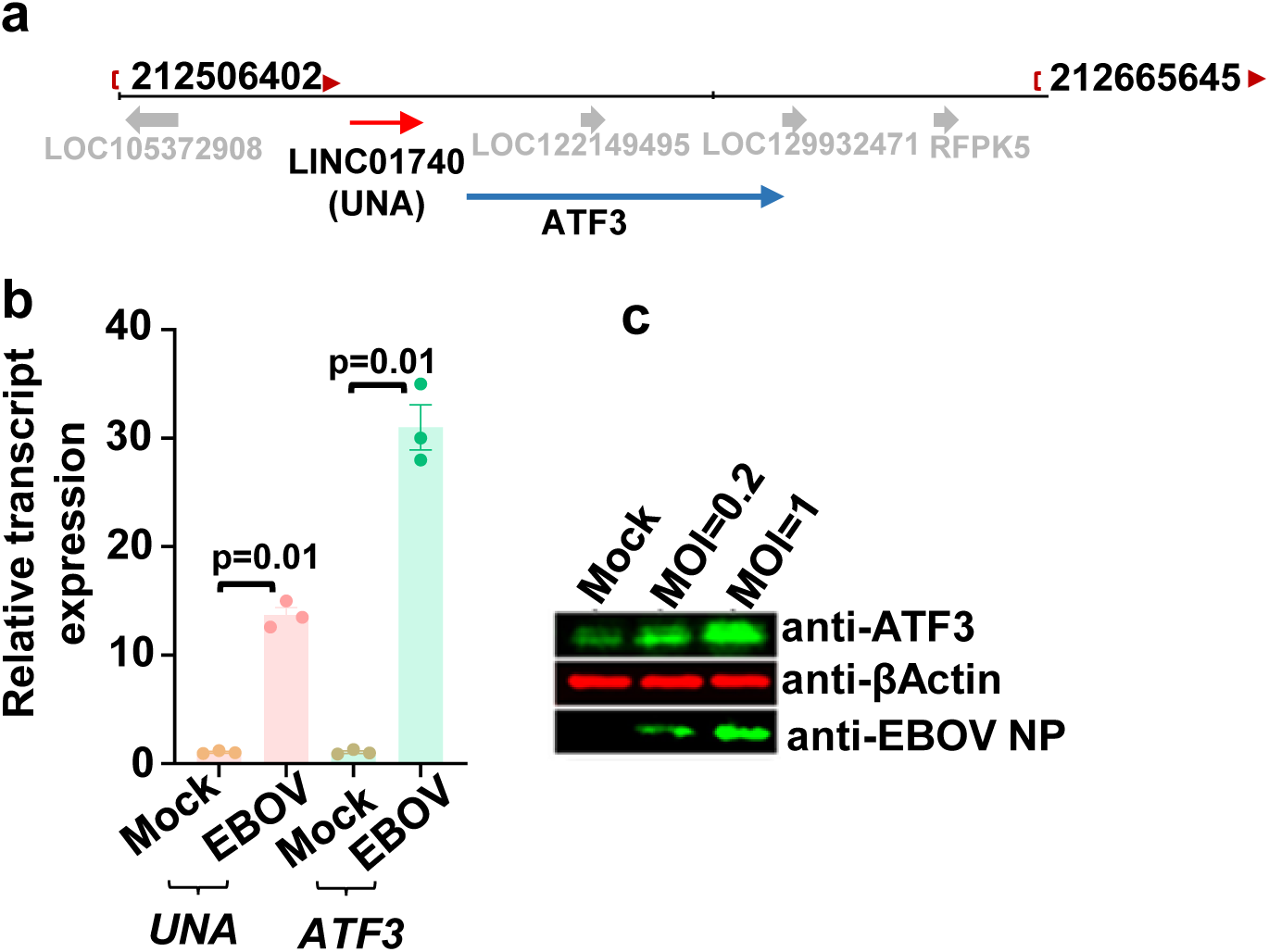
LINC01740 (UNA) and the neighboring protein-coding gene ATF3 are significantly upregulated in EBOV-infected human macrophages. **(a)** Genomic localization of LINC01740 (UNA) and ATF3 on human chromosome 1. **(b)** mRNA expression levels of UNA and ATF3 are significantly increased at 24 h post–EBOV infection compared with mock-infected macrophages. Data are presented as mean ± SEM. Statistical comparisons were made using Student’s t-test, and two-tailed p-values are indicated. **(c)** ATF3 protein expression is increased in EBOV-infected human macrophages. Cells were infected with EBOV at a multiplicity of infection (MOI) of 0.2 or 1, and cell lysates were collected at 24 h post-infection for immunoblot analysis of ATF3 and EBOV nucleoprotein (NP).

Although the UNA (LINC01740) transcript did not contain any open reading frames (ORF) longer than 20 codons, studies have suggested that several lncRNAs may encode short peptides [48]. To test this possibility, we analyzed the UNA transcript sequence using the coding potential calculator 2 (CPC2) [49]. CPC2 uses the Fickett score to distinguish between coding and non-coding nucleotide sequences. A higher Fickett score indicates a higher likelihood of coding potential, while a lower score suggests a higher likelihood of a non-coding sequence. UNA Sequence (LINC01740, ENST00000423842.2) yielded a Fickett score of 0.31252, a complete putative ORF of 25 amino acids, and a pI of 6.50531005859, which together classify it as a noncoding sequence with a coding probability of 0.00481494

### UNA enhances the expression of ATF3

LncRNAs can regulate the expression of neighboring genes. Given that the ATF3 and UNA genes are within 10 kb of each other (Fig. 2a), we tested whether UNA regulates ATF3 expression. We designed antisense oligonucleotides (ASOs) targeting the UNA transcript to knock down UNA (UNA-KD). We differentiated THP-1 cells into macrophage-like cells using phorbol 12-myristate 13-acetate (PMA) stimulation and transfected cells with negative control ASO (NC) or UNA-targeting ASO (UNA-KD). Our data show a significant reduction in ATF3 mRNA and protein levels in UNA-KD cells compared with NC-transfected THP-1 macrophages (Fig. 3a, b, c). Similarly, we observed a significant decrease in ATF3 mRNA levels in UNA-KD vs. NC-transfected cells in other cell types, including human kidney 293 T-derived LentiX cells and epithelial cells (HeLa) (Fig. S4). Pol II Chromatin Immunoprecipitation (ChIP) followed by qPCR (ChIP-qPCR) shows significantly reduced Pol II occupancy at the ATF3 promoter in UNA-KD compared with NC THP-1 macrophages (Fig. 4), supporting a role for UNA in promoting ATF3 transcription. We also investigated whether UNA regulates post-transcriptional decay of ATF3 mRNA. Knockdown of UNA expression did not affect ATF3 mRNA decay (Supplementary Fig. S5). Taken together, these data indicate that UNA enhances ATF3 transcription.

**Figure 3.**
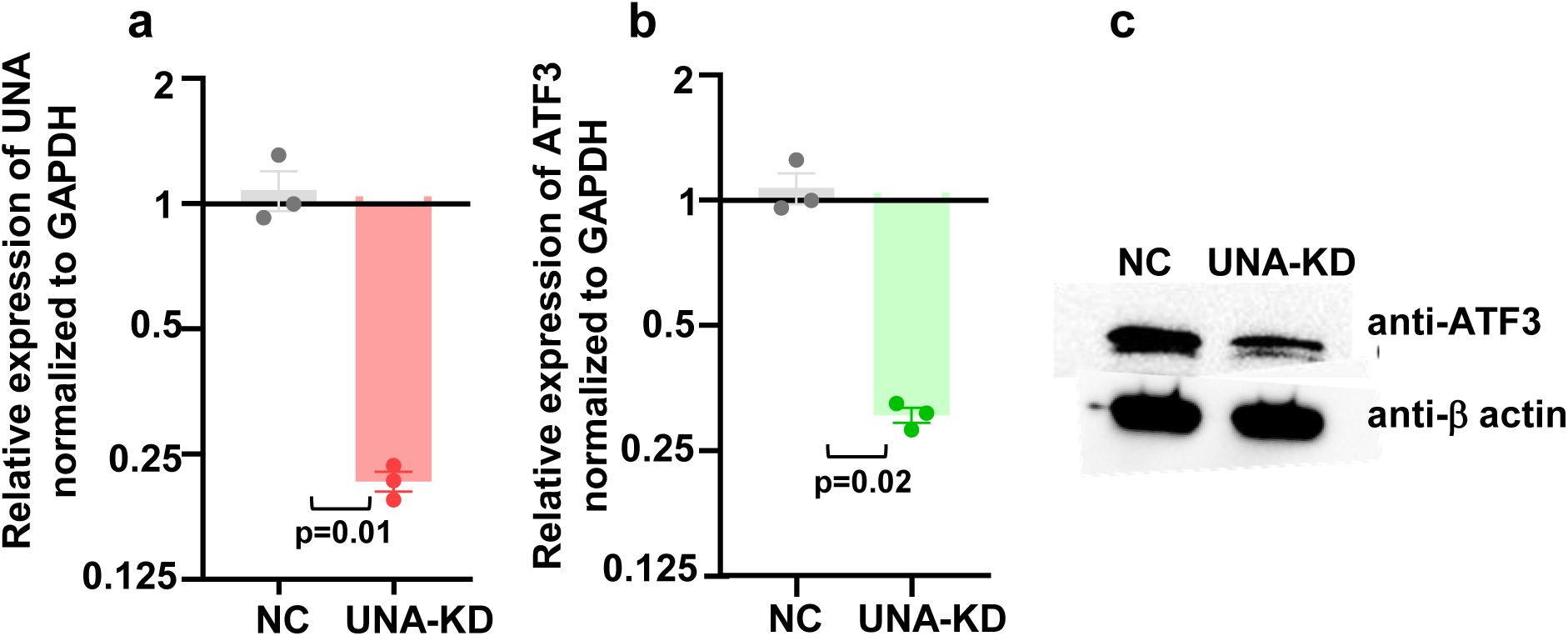
UNA knockdown reduces ATF3 mRNA and protein expression in PMA-differentiated THP-1 cells. Cells were transfected with an antisense oligonucleotide (ASO) targeting UNA (UNA-KD) or a negative control ASO (NC). **(a)** UNA transcript levels were significantly reduced in UNA-KD cells than in NC-transfected cells. ATF3 mRNA **(b)** and protein expression **(c)** were also significantly decreased in UNA-KD cells than in controls. Data are presented as mean ± SEM. Statistical comparisons were made using Student’s t-test, and two-tailed p-values are indicated

**Figure 4.**
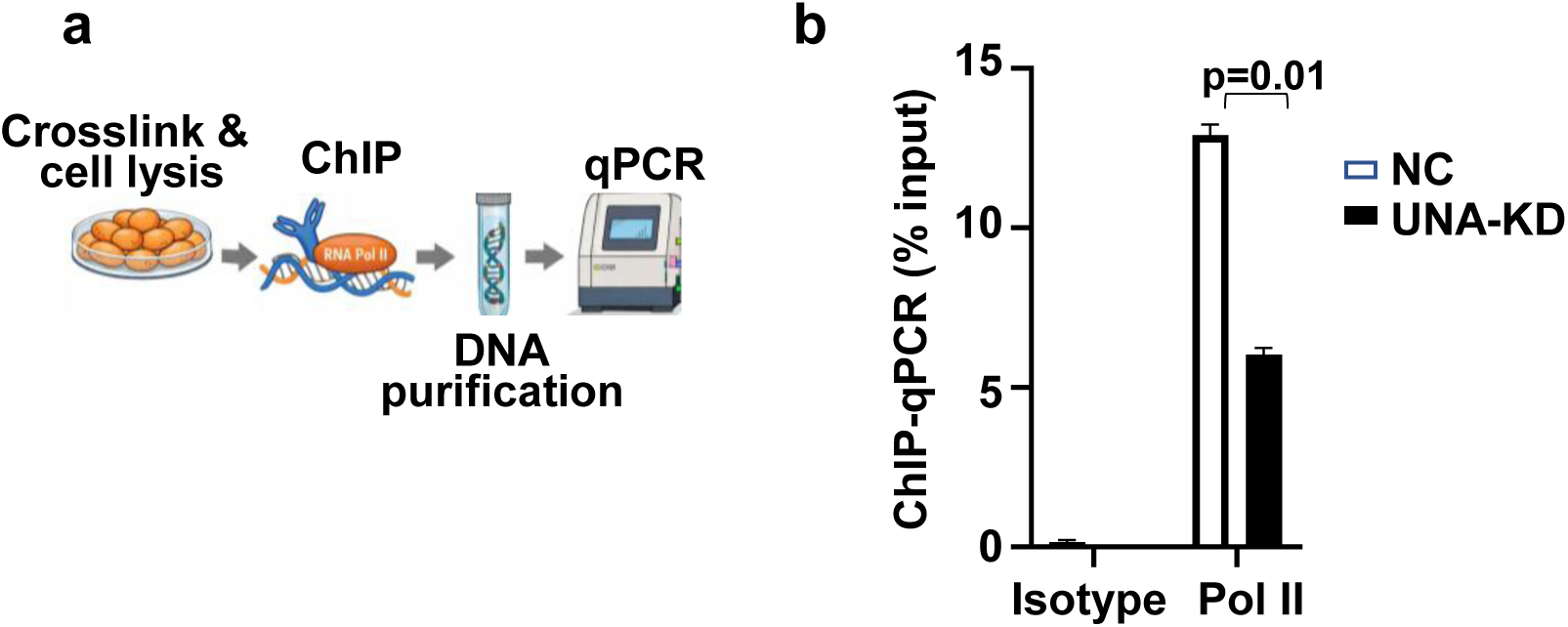
UNA silencing inhibits ATF3 transcription. **(a)** THP-1 cells were differentiated with PMA and then transfected with antisense oligos (ASOs) to knock down UNA (UNA-KD) or a negative control (NC) ASO. 24 h post-transfection, the cells were crosslinked and lysed. We then performed Chromatin Immunoprecipitation (ChIP) using an RNA polymerase II (Pol II) antibody or an isotype control. We measured enrichment of the ATF3 promoter in Pol II-associated chromatin using specific primers. (**b)** ChIP-qPCR analysis of RNA Pol II in UNA-knockdown and control cells (UNA-KD vs. NC). The mean ± SE is shown as horizontal and vertical bars for each group. Student’s t-test was used for statistical comparison, and a two-tailed p-value is indicated.

### ATF3 enhances viral replication in human macrophages by suppressing the antiviral response

To determine the impact of ATF3 on EBOV virus replication, we used Inducible CRISPR/Cas13d-mediated RNA Degradation (iCRISPRd) for the transient knockdown of ATF3 expression. Briefly, we transduced THP-1 cells with lentiviral particles encoding doxycycline-inducible CRISPR/Cas13d and a blasticidin resistance gene. We selected cells with stable inducible CRISPR/Cas13d integration in media containing blasticidin for several days. Selected cells were then transduced with lentiviral particles encoding an ATF3-targeting guide RNA (gATF3) or a non-targeting guide RNA (gNT) and a puromycin resistance gene. We selected cells in media with blasticidin and puromycin for several days. gATF3 and gNT cells were differentiated into macrophage-like cells, and CRISPR-Cas13d expression was induced with doxycycline (Fig. 5a). We measured ATF3 knockdown in gATF3 vs. gNT cells by qPCR and immunoblot (Fig. 5b, c). We infected gATF3 and gNT cells with EBOV encoding GFP (EBOV-GFP) at an MOI of 0.2 for 24 h, and then the numbers of infected cells and nuclei were determined by microscopy. Relative EBOV-GFP expression was significantly reduced in gATF3 vs. gNT cells (Fig. 5d). We confirmed the impact of ATF3 inhibition on EBOV replication across distinct cell types and using a different method to inhibit ATF3 expression. We observed a similar effect of shRNA-mediated ATF3 knockdown on viral replication in liver (Huh) or epithelial (HeLa) cell lines. EBOV replication was significantly downregulated in ATF3-targeting vs. negative control shRNA (ATF3-sh vs. NC-sh; Supplementary Fig. S6).

**Figure 5.**
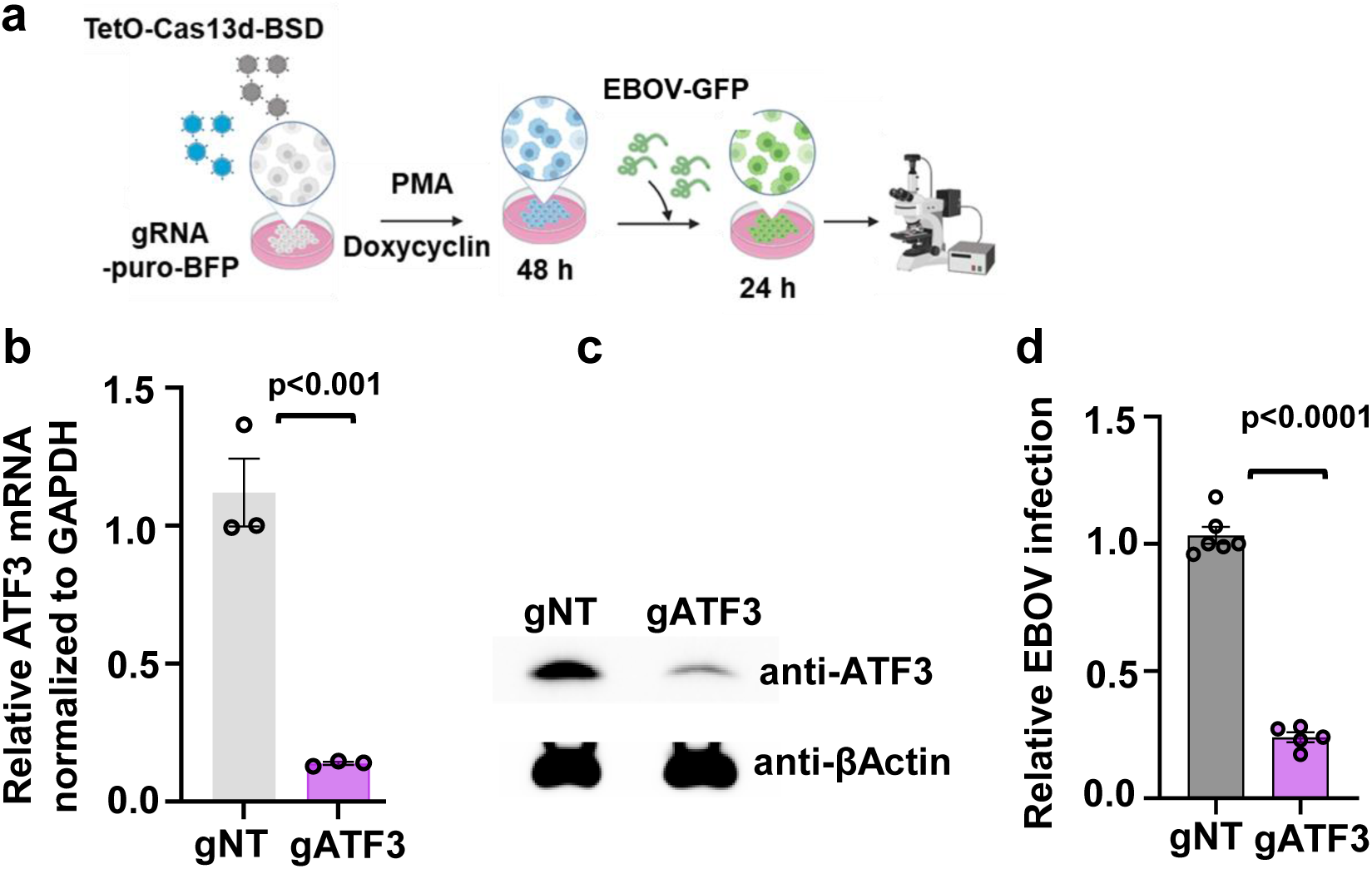
ATF3 depletion reduces EBOV infection in THP-1-derived macrophages. **(a)** PMA-differentiated THP-1 cells expressing CRISPR/Cas13d and either a nontargeting guide RNA (gNT) or an ATF3-targeting guide RNA (gATF3) were infected with EBOV-GFP (MOI, 0.2). At 24 h post-infection, EBOV-GFP-positive cells and total nuclei were quantified by automated fluorescence microscopy. **(b and c)** Knockdown of ATF3 was confirmed by RT-qPCR **(b)** and immunoblotting **(c). (d)** EBOV-GFP infection was significantly reduced in gATF3 cells compared with gNT controls. . Data are presented as mean ± SEM. Statistical comparisons were made using Student’s t-test, and two-tailed p-values are indicated.

ATF3 regulates a broad transcriptional network encompassing inflammation, autophagy, and apoptosis (reviewed in [50, 51]). To define its role in antiviral immunity in EBOV infection, we performed RNA-seq on EBOV-infected THP-1 macrophages following ATF3 knockdown (gATF3) or control treatment (gNT). Gene set enrichment analyses (GSEA) showed that the loss of ATF3 markedly enhanced IFN-I-mediated antiviral and inflammatory pathways, indicating that ATF3 acts as a negative regulator of innate immune activation. ATF3 depletion activated interconnected modules governing macrophage activation and antiviral defense. Expression of proinflammatory cytokines was significantly increased in EBOV-infected THP-1 macrophages with ATF3 knockdown (gATF3) than in control cells (gNT), consistent with enhanced macrophage-driven inflammatory signaling. The analyses also reveal prominent Toll-like receptor (TLR) signaling and downstream induction of IFN-I pathways, which promote pathogen sensing, phagocytic activation, and antigen presentation. In ATF3-knockdown macrophages, a robust interferon signaling module, including IFN and known type I interferon-stimulated genes (ISGs), was evident, reflecting a coordinated transcriptional program that may augment antiviral gene expression and restrict viral replication. ISGs with known antiviral function (indicated in Fig. 6a, Supplementary Table S2) are significantly upregulated in gATF3 cells, indicating that ATF3 inhibition enhances antiviral transcriptional programs that restrict EBOV replication. In addition, ISGs known to block EBOV entry, such as ISG15 [52] and IFITM1 [53, 54], are upregulated in ATF3-silenced cells. Concurrently, activation of the TNF, TGF, and JAK-STAT pathways amplifies inflammatory crosstalk (Fig. 6b). Many differentially expressed genes (DEGs) are transcriptional regulators and direct targets of ATF3, suggesting that ATF3 regulates a hierarchical gene network that governs macrophage function during EBOV infection.

**Figure 6.**
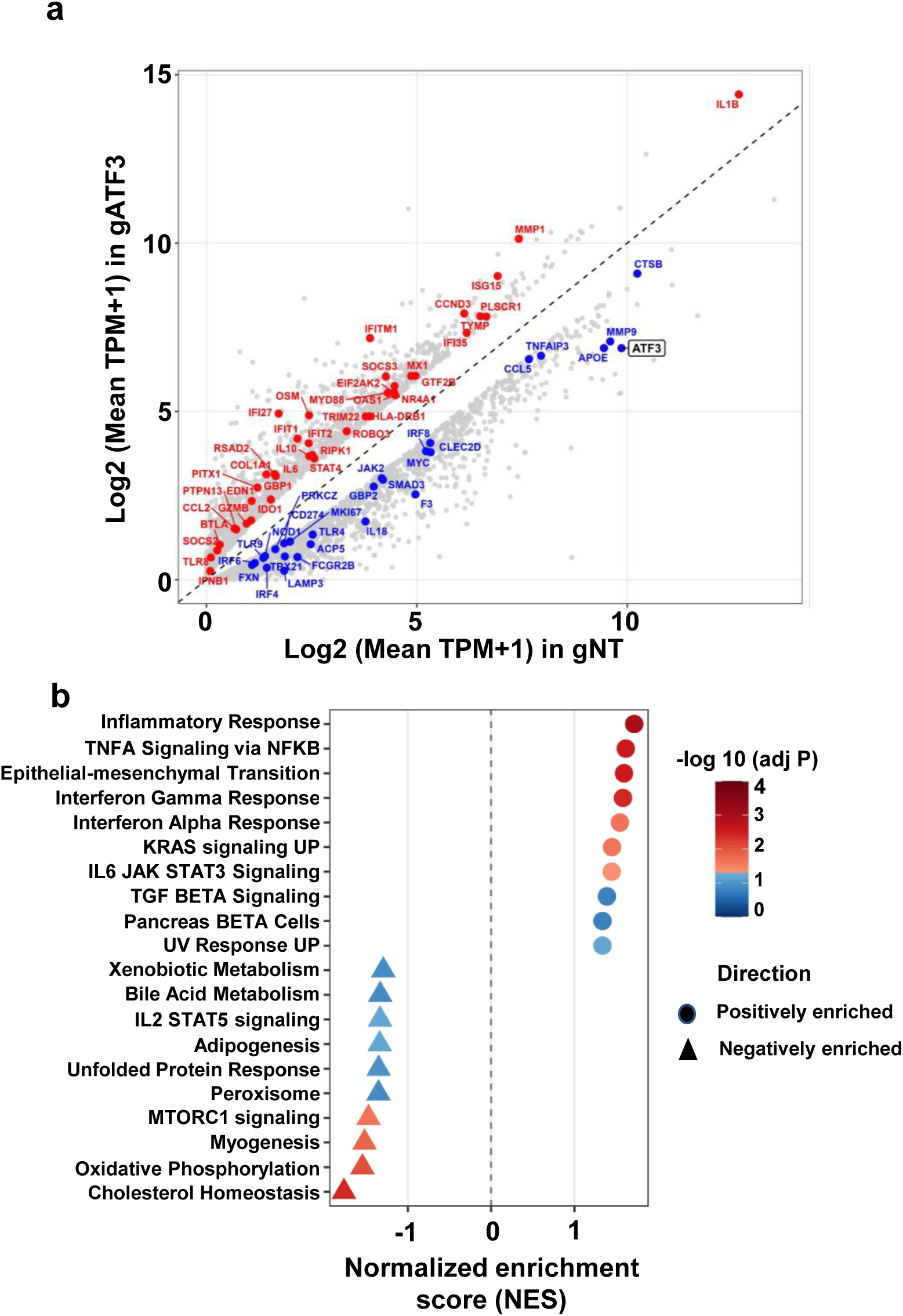
ATF3 depletion enhances interferon-associated gene expression in EBOV-infected THP-1-derived macrophages. **(a)** RNA sequencing was performed on EBOV-infected THP-1-derived macrophages expressing an ATF3-targeting guide RNA (gATF3) or a nontargeting control guide RNA (gNT). The scatter plot (log2[TPM + 1]) gene expression values in gATF3 (y-axis) versus gNT (x-axis). Each point represents a differentially expressed gene. Differentially expressed interferon-associated genes are labeled, with genes upregulated in gATF3 and gNT shown in red and blue, respectively. **(b)** Hallmark gene set enrichment analysis (GSEA) of differentially expressed genes. Pathways are ranked by normalized enrichment score (NES). Dot size represents gene set size, and color indicates the –log10 (adj p-value). Positive NES values indicate enrichment in gATF3 cells, whereas negative NES values indicate enrichment in gNT cells.

Collectively, these findings demonstrate that EBOV-induced ATF3 expression suppresses key IFN-I and inflammatory responses in macrophages, thereby delaying protective antiviral responses and promoting viral replication.

## Discussion

Ebola virus (EBOV) infection triggers profound dysregulation of host immunity, including impaired IFN-I responses and hyperinflammation, both of which contribute to the pathogenesis of EVD [55–60]. Macrophages are primary targets of EBOV, and viral replication extensively alters their transcriptional programs. Although lncRNAs are increasingly recognized as regulators of immune responses and viral pathogenesis [9, 17–25], their roles during EBOV infection remain poorly defined. Here, we identify a lncRNA-transcription factor axis in primary human macrophages that modulates antiviral immunity and facilitates viral replication.

Transcriptomic profiling of EBOV-infected monocyte-derived macrophages revealed induction of hundreds of EVILs in the genomic proximity of protein-coding genes regulating cytokine signaling, transcriptional regulation, apoptosis, and other cell-signaling pathways relevant to EVD. Among the top EVILs, UNA (LINC01740), located upstream of ATF3, was strongly upregulated and co-expressed with ATF3. ASO-mediated knockdown of UNA reduced ATF3 mRNA and protein levels, confirming that UNA positively regulates ATF3 transcription.

ATF3 is a member of the activating transcription factor/cAMP response element–binding (ATF/CREB) family and is rapidly induced by cellular stress, Toll-like receptor signaling, and viral infection [61, 62]. ATF3 is shown to have context-dependent antiviral or proviral functions in several viral infections, including hepatitis B virus [63, 64], Herpes simplex virus [65], human papillomavirus [66, 67], Japanese encephalitis virus [62], Zika virus [68], Coxsackievirus [69], Dengue virus [70], and HIV [71]. ATF3 functions as both a transcriptional activator and a repressor, depending on its dimerization partners and promoter context. In macrophages, ATF3 acts as a negative feedback regulator that attenuates excessive inflammatory signaling, in part by repressing NF-κB–dependent transcription [72–74] and inhibiting IFNβ promoter activity [61]. Consistent with these roles, silencing ATF3 in EBOV-infected macrophages restored IFN-I signaling and increased expression of interferon-stimulated genes (ISGs) and inflammatory mediators, indicating that ATF3 serves as a transcriptional checkpoint that dampens antiviral immunity. Mechanistically, ATF3 may repress IFN-I-responsive genes through promoter occupancy, recruitment of chromatin-modifying complexes, or interference with transcription factors such as IRF3 and STAT1.

These findings support a model in which ATF3 facilitates EBOV-mediated suppression of IFN-I–dependent antiviral responses, thereby promoting viral replication. Consistent with this model, ATF3 depletion significantly reduced EBOV replication across cell types, underscoring its role in facilitating EBOV immune evasion. EBOV-induced LINC01740 contributes to this process by enhancing ATF3 transcription, reinforcing ATF3-mediated negative feedback within antiviral gene networks. Elevated ATF3 activity delays the establishment of an effective antiviral state, enabling EBOV to evade host innate immune defenses more efficiently.

The LINC01740–ATF3 axis exemplifies a broader paradigm in viral immune modulation. Many viruses exploit host transcription factors to recalibrate inflammatory thresholds. EBOV leverages this lncRNA-dependent mechanism to reinforce suppression of IFN-I signaling, demonstrating that infection-responsive lncRNAs can be active determinants of host defense rather than passive byproducts of infection. EBOV has evolved multiple strategies to suppress and delay IFN-I induction and the expression of interferon-stimulated antiviral genes, enabling viral replication and immune dysregulation. Viral proteins such as VP35 and VP24 potently antagonize IFN-I expression and signaling [55–60]. Impaired IFN-I responses in macrophages and dendritic cells contribute to the “immune paralysis” characteristic of Ebola virus disease, allowing the virus to disseminate systemically while driving hyperinflammation and tissue damage. The survivors of EBOV infection show significant upregulation in IFN signaling as indicated by higher expression of ISGs (IFIT1, IFI6, IFIT3, ISG15, OAS1, IFITM3*, and* MX1) [75]. Ebola virus survivors show earlier IFN-I responses and inflammatory cytokine (IL-1β, IL-6, TNFα) expression than the non-survivors [76, 77]. Notably, several of these ISGs and inflammatory cytokines are regulated by ATF3 expression in our dataset (Fig. 6). Notably, early ATF3 induction after Ad26.ZEBOV vaccination correlates with enhanced antibody responses [78], underscoring the context-dependent immunomodulatory function of ATF3.

Several limitations warrant consideration. First, although LINC01740 enhances ATF3 transcription, the precise molecular mechanism remains undefined. Chromatin accessibility mapping, histone modification profiling, and chromosome conformation analyses will be required to distinguish among enhancer-like activity, RNA-dependent scaffolding, and transcription-coupled chromatin remodeling. Second, these findings derive from in vitro macrophage models; in vivo studies are needed to establish the contribution of the LINC01740–ATF3 axis to systemic EBOV pathogenesis. Third, although ATF3 depletion reduced viral replication, the downstream effectors responsible for this phenotype remain to be fully delineated.

In conclusion, we describe a previously unrecognized LINC01740-ATF3-IFN-I regulatory axis in EBOV-infected macrophages. LINC01740 enhances ATF3 transcription, which in turn suppresses IFN-I–mediated antiviral responses, creating a cellular environment permissive for viral replication. These findings align with emerging evidence that host lncRNAs are dynamically regulated during EBOV infection and may coordinate antiviral responses. For example, single-cell profiling of rhesus macaque immune cells revealed differential lncRNA expression during EBOV infection and frequent coexpression with protein-coding immune regulators, underscoring the broad relevance of infection-responsive noncoding RNAs in filovirus pathogenesis [26]. Recent transcriptomic studies of Ebola virus vaccination further suggest that lncRNAs are integral components of protective immunity, with conserved lncRNA expression associated with interferon signaling, T-cell differentiation, and antibody production against EBOV [79]. Together, these observations support a model in which infection-responsive lncRNAs act as transcriptional rheostats that fine-tune host immune responses. In this context, EBOV appears to co-opt LINC01740 to enhance ATF3 expression and suppress IFN-I signaling, potentially contributing to the delayed interferon responses characteristic of severe Ebola virus disease. These findings highlight the broader importance of lncRNAs as regulators of infection outcomes.

## Supporting information

Supplementary Table 2. Significantly Differentially Expressed Genes in EBOV-Infected THP-1 Macrophages Following ATF3 Knockdown

## Acknowledgements

The project was supported in parts by NIH/NIAID (AI157850; S. K), Texas Biomed Forum grants (S.K, O.S), Texas Biomed Forum post-doctoral grant (T.G.), Cowles post-doctoral fellowship (A.D.), and Institutional funds from the Texas Biomedical Research Institute (S.K. and O.S)

## Author Contributions

O.S. and S.K. designed the study. O.S., T.G., A.G., M.D.L., A.D., S. J., A.T., H.N., and S.K. designed the experiments, performed the experiments, analyzed the data, and interpreted the results. S.K. directed the study and wrote the manuscript with T.G. All authors provided intellectual input and edited the manuscript.

## Methods

### Samples

Primary human macrophages were differentiated from buffy coats obtained from 9 donors at the South Texas Blood and Tissue Center using a published protocol [80–82]. Mononuclear lymphocytes were isolated using LeucoSep tubes (Fisher Scientific), resuspended in Iscove’s modified Dulbecco’s medium (IMDM, Fisher Scientific), and plated in 100-mm plates for 1 hour to adhere. Unattached cells were washed off, and attached monocytes were differentiated into macrophages for 7 days in IMDM supplemented with 2% heat-inactivated human serum (Corning), 100 U/mL penicillin, 100 µg/mL streptomycin, nonessential amino acids, 50 mM 2-mercaptoethanol, and 800 U/mL human macrophage colony-stimulating factor. Adherent monocytes were washed, and the medium was replaced on days 2 and 6. All cells were incubated at 37°C in a humidified incubator with 5% CO2.

### EBOV infection

Wild-type EBOV (Mayinga, NCBI accession number AF086833) and replication-competent Ebola virus expressing green fluorescent protein (EBOV-GFP, NCBI accession number KF990213.1) were handled in a maximum-containment (BSL-4) laboratory at Texas Biomedical Research Institute (San Antonio, TX, USA) in accordance with institutional standard operating procedures approved by the Biohazard and Safety Committee and the Recombinant DNA Committee. Virus stocks were generated by propagating the virus in Vero cells maintained in DMEM (Corning) supplemented with fetal bovine serum for 7 days. Culture supernatants were clarified to remove cellular debris, layered onto a 20% sucrose cushion in phosphate-buffered saline (PBS), and centrifuged at 28,000 rpm for 2 hours at 4°C. Viral pellets were resuspended in PBS and stored at −80°C until use. Viral titers were determined by infecting Vero cells with serial dilutions of virus for 24 hours and quantifying fluorescent focus–forming units per milliliter (FFU/mL).

THP-1 cells were seeded into 24-well plates at 10^6 cells/well and left uninfected or incubated with EBOV at a multiplicity of infection (MOI) of 0.2, 0.5, or 1 (as titrated in Vero cells). After 1 h, cells were washed twice with PBS and incubated in fresh medium for 24 h. Cells were fixed and stained with Hoechst dye (Thermo Fisher) to label nuclei, and images were acquired using an automated Nikon Ti-Eclipse microscope (Nikon, Tokyo, Japan) as described [83]. The number of cell nuclei and infected (GFP-positive) cells was counted using CellProfiler v4.2.3 (Broad Institute).

### RNA extraction and quantitative PCR (qPCR)

Total RNA was extracted from cell lines or primary macrophages using the RNeasy Universal kit (Qiagen) or the Direct-zol™ RNA MiniPrep (Zymo Research). Each sample was treated with on-column DNase to remove genomic DNA. RNA was quantified using a NanoDrop spectrophotometer (Thermo Fisher). mRNA expression levels were measured by qPCR. Briefly, reverse transcription was performed on 1000 ng of total RNA using the High-Capacity RNA-to-cDNA Kit (Applied Biosystems) in a 10 µl reaction volume. Specific protein-coding regions (ATF3, GAPDH) and non-coding regions (LINC01740) were amplified by SYBR green qPCR on QuantStudio 5 instruments (Applied Biosystems). Each qPCR reaction contained 5 µl of Power SYBR Green PCR Master Mix (Applied Biosystems), 200 nM primers specific for the gene of interest or the housekeeping gene (GAPDH), and 4 µl of cDNA (1:4 dilution), for a total volume of 10 µl. The genes were amplified under the following conditions: 50°C for 2 minutes, 95°C for 10 minutes, and 40 cycles of 95°C for 15 seconds and 60°C for 1 minute. Primer specificity was assessed by melt curve analysis, with a dissociation step performed as per the qPCR protocol. Gene expression levels were normalized to a housekeeping gene (GAPDH) using the 2(-Delta Delta CT) method [84]. Oligonucleotide primer sequences are provided in the list of primers and probes in Supplementary Table S1.

### ATF3 mRNA decay

ATF3 mRNA decay was assessed by qPCR. THP-1 cells were differentiated with PMA (Millipore Sigma) for 72 hours, then transfected with UNA-targeting or control ASOs. Twenty-four hours post-transfection, cells were treated with Actinomycin D or DMSO for 0, 60, or 120 minutes. Cells were trypsinized, washed with PBS, and stored in TRI reagent until RNA extraction. Total RNA was isolated, quantified, and used for cDNA synthesis and qPCR analysis of UNA, ATF3, and GAPDH (primer sequences are provided in Supplementary Table S1).

### Plasmids and lentiviruses CRISPR/Cas13d

CRISPR/Cas13 lentiviral vector Cas13d-TetO-BlastR (pLentiRNACRISPR_007-TetO-NLSp-RfxCas13d-NLS-WPRE-EFS-rTA3-2A-Blast was a gift from Neville Sanjana; Addgene plasmid #138149); gRNA expression vectors (pXR003 was a gift from Patrick Hsu; Addgene #109054); Vesicular Stomatitis Virus glycoprotein (VSV-G) envelope expression vector (pMD2.G was a gift from Didier Trono; Addgene #12259); lentiviral packaging plasmid (psPax2 was a gift from Didier Trono; Addgene #12260); and gRNA cloning plasmid [pKLV2-U6gRNA5(BbsI)-PGKpuro2ABFP-W was a gift from Kosuke Yusa; Addgene #67974] were obtained from Addgene (Watertown, MA, USA).

### Construction of the pKLV2-U6-CasRx-(gRNA)-PGKpuro2ABFP (pKLV2-XR3) vector

The pKLV2-U6-CasRx-(gRNA)-PGKpuro2ABFP vector was constructed by restriction enzyme digestion and ligation. The gRNA cassette was PCR-amplified using Q5® High-Fidelity DNA polymerase (New England Biolabs, Ipswich, MA, USA) from the pXR003 plasmid (Addgene #109053) to include MluI and BamHI restriction sites (PCR primers in Supplementary Table S1). One microgram of this PCR product and pKLV-U6gRNA (BbsI)-PGKpuro2ABFP (Addgene #67974) were digested with MluI and BamHI (New England Biolabs, Ipswich, MA, USA). The digested products were purified, ligated, and transformed into E. coli Stbl competent cells (New England Biolabs, Ipswich, MA, USA). Colonies were screened by PCR, and positive clones were grown. The vector sequence was further confirmed by Sanger sequencing as described above. Sanger sequencing primers are listed in Supplementary Table S1.

### gRNA cloning

10 µM of each gRNA primer pair was phosphorylated and annealed in a 10 µL reaction containing 1 µL of 10× T4 DNA Ligase Buffer, 0.5 µL of T4 polynucleotide kinase (New England Biolabs, Ipswich, MA, USA), and distilled water. Phosphorylation was performed at 37 °C for 30 min. The primer pairs were annealed by heating the reaction to 95 °C for 5 min and then cooling to 25 °C at 5 °C/min. The phosphorylated and annealed gRNAs were cloned into pKLV2-XR3 using a Golden Gate cloning reaction. Each reaction contained 1 µM annealed crRNA guide oligonucleotides, 25 ng of pKLV2-XR3, 0.5 µL of BbsI (10 U/µL), 0.5 µL of T4 DNA Ligase (New England Biolabs, Ipswich, MA, USA) (400 U/µL), 1 µL of 10× T4 DNA Ligase Buffer, and distilled water, for a total of 10 µL. The Golden Gate amplification was carried out for 30 cycles of 37 °C for 5 min and 23 °C for 5 min. The Golden Gate mix was transformed into E. coli Stbl competent cells (New England Biolabs, Ipswich, MA, USA), and the gRNA sequences are listed in Supplementary Table S1.

### Cell lines and transfection

THP-1 cells were maintained in RPMI 1640 medium (Gibco) supplemented with 10% heat-inactivated fetal bovine serum (FBS; Atlanta Biologicals). 293 T-derived LentiX, HeLa, and Huh cells were maintained in DMEM (Gibco) supplemented with 10% heat-inactivated FBS (Atlanta Biologicals). THP-1 cells were differentiated with PMA for 72 hours at 1 million cells/mL in a 12-well plate. Differentiated THP-1 cells were then transfected with Antisense Oligonucleotides (ASO) using Lipofectamine^TM^ 2000 (Invitrogen).

### Lentivirus transduction and cell line selection

CRISPR/Cas13d or pKLV2-RX3 gRNA lentiviral particles were produced by co-transfecting Lenti-X^TM^ cells with plasmids encoding Cas13d-TetO-BlastR or gRNAs cloned into pKLV2-XR3, along with VSV-G envelope (pMD2.G; Addgene, Cambridge, MA, USA) and packaging (psPAX2; Addgene, Cambridge, MA, USA) plasmids, using Lipofectamine 3000 (Thermofisher Scientific) to generate CRISPR/Cas13d and gRNA-encoding lentiviral particles, respectively. Lentiviral supernatant was harvested 72 h after transfection, centrifuged at 500g for 10 min at 4 °C to remove cellular debris, and concentrated 10-fold using Lenti-X concentrator (Takara Bio, Mountain View, CA, USA). To obtain cells with stable Cas13d expression, THP1 cells were transduced with Cas13d-TetO-BlastR lentivirus in growth media containing 8 µg/mL polybrene. 72 hours after transduction, the growth medium was replaced with cRPMI containing Blasticidine (6 µg/mL), and the cells were selected for several cycles. Cells with stable Cas13d were then transduced with lentiviral particles encoding an ATF3-targeting gRNA (gATF3) or a negative control gRNA (gNT). After 72 hours post-transduction, cells were selected in media containing Blasticidine and puromycin (5 µg/mL) for several cycles. Selected cells were differentiated using phorbol 12-myristate 13-acetate (PMA) for 24 hours and treated with doxycycline for 48 hours. Reductions in ATF3 mRNA and protein levels were measured by qPCR and Immunoblotting, respectively.

### Immunoblot

THP-1 cells were lysed in buffer containing 20 mM Tris-HCl (pH 7.4), 150 mM NaCl, 1% NP-40, and 5 mM EDTA, supplemented with 1x Protease Inhibitor Cocktail (Promega). Clarified lysates were resolved by 4%-20% SDS-PAGE (Invitrogen) and transferred to a 0.2 μM PVDF membrane using Turbo-Blot (Bio-Rad). Membranes were blocked with EveryBlot blocking buffer (Bio-Rad) for 1 hr and probed overnight with primary antibodies in blocking buffer. The antibodies used in Western blots were ATF3 and β-actin (Cell Signaling Technology). Membranes were probed with horseradish peroxidase-conjugated anti-rabbit and anti-mouse secondary antibodies (Bio-Rad; cat. 172-1011 and 170-6515). Western blots were developed using ECL chemiluminescent substrate (Pierce).

Primary macrophages infected with EBOV were lysed in RIPA buffer (50 mM Tris-Cl [pH 7.5], 150 mM NaCl, 1% Triton X-100, and 0.1% SDS). The lysates were kept on ice for 15 minutes and then clarified by centrifugation. The lysates were incubated with SDS-PAGE sample buffer at 100°C for 10 minutes, and proteins were resolved on a 4%–20% SDS-PAGE gradient gel (Bio-Rad Laboratories, Hercules, CA). The samples were transferred onto iBlot nitrocellulose membranes (Life Technologies, Carlsbad, CA) and blocked for 1 h at room temperature with blocking buffer (LI-COR Biosciences, Lincoln, NE). The blots were incubated with primary antibodies (ATF3 and EBOV NP) at 4°C overnight, followed by incubation with anti-rabbit IRDye 800CW and anti-mouse IRDye 680LT (LI-COR Biosciences, Lincoln, NE) for 1 h at room temperature. Protein bands were visualized using Odyssey software (LI-COR Biosciences, Lincoln, NE).

### lncRNA microarray

RNA was extracted using the RNeasy Universal Plus kit (QIAGEN). RNA concentration was measured with a NanoDrop spectrophotometer, and RNA integrity was assessed using a Fragment Analyzer (Agilent). LncRNA profiles were obtained using the Arraystar Human lncRNA Microarray (V3.0, Agilent Technologies, Santa Clara, CA, USA), designed for global profiling of 30,586 human lncRNAs and 26,109 protein-coding transcripts. After sample labeling and array hybridization, raw data were extracted using the Agilent Feature Extraction software (version 11.0.1.1). Quantile normalization and subsequent data processing were performed using the GeneSpring GX v12.1 software package (Agilent Technologies). After quantile normalization, lncRNAs and mRNAs with signal quality flags of “Present” or “Marginal” in at least 25% of the samples were retained for downstream analysis. Fold change (FC) was calculated from log₂-transformed normalized expression values using the formula FC = 2^(|norm1 − norm0|), where norm1 and norm0 represent the mean normalized expression values of the two comparison groups. Differential expression analysis was assessed using a two-tailed, homoscedastic Student’s t-test on normalized expression intensities, followed by Benjamini–Hochberg false discovery rate (FDR) correction for multiple testing. lncRNAs/mRNAs with an FDR-adjusted p-value < 0.05 and an absolute log₂ fold change ≥ |1.0| were considered significantly differentially expressed.

### RNA sequencing

Cell lysates were collected in TRIzol® Reagent (Ambion, Inc.). RNA was isolated using the Direct-zol™ RNA MiniPrep (Zymo Research) according to the manufacturer’s instructions, including DNase treatment. RNA concentration and purity were assessed using a Nanodrop spectrophotometer (Thermo Fisher Scientific), and samples were shipped on dry ice. Before RNA sequencing, RNA quality was assessed using the Fragment Analyzer (Agilent). Samples with an RNA integrity number (RIN) above 7 were used for RNA-Seq. RNA-Seq libraries were prepared from 500 ng of total RNA using the NEB Next directional RNA library preparation kit with poly(A) enrichment module (Ipswich, MA). RNA sequencing was performed at the Genome Sequencing Facility (GSF) at UT Health San Antonio on the NovaSeq 6000 System (Illumina) with 100-bp paired-end reads.

### Analysis of differentially expressed genes

Raw paired-end RNA-seq reads were initially assessed with FastQC v0.12.0 (Babraham Bioinformatics - FastQC: A Quality Control tool for High Throughput Sequence Data). Low-quality reads and adapter sequences were trimmed using Trim Galore v0.6.10 (Babraham Bioinformatics - Trim Galore!). Transcript abundance was quantified via pseudo-alignment to the human transcriptome (Homo sapiens GRCh38.p14 cDNA) using kallisto (version 0.51.1) [85]. Gene-level summaries were generated with the tximport R package (version 1.36.1) [86]. Only genes annotated as protein-coding based on GENCODE v49 gene biotype annotations were retained for downstream differential expression and enrichment analyses. Differential gene expression analysis between experimental groups was performed using DESeq2 (v1.48.1). Significantly differentially expressed genes were identified using a log2 fold change ≥ |1.0| and FDR-adjusted p-values (Benjamini-Hochberg method) < 0.05.

To visualize biologically relevant antiviral responses, a curated set of interferon-associated genes was obtained from the Harmonizome database[87] using the GeneRIF Biological Term Annotations dataset and querying with the term “interferon.” Gene-level TPM values generated with tximport were averaged across biological replicates for each condition and then transformed as log2(TPM + 1). Fast Gene Set Enrichment Analysis (fGSEA) was performed with 1000 permutations, using preranked differential expression statistics based on the Wald test statistic (“stat” column from DESeq2 results). Hallmark gene sets for Homo sapiens were sourced from the Molecular Signatures Database (Human MSigDB v2026.1.Hs). Enrichment significance was assessed through normalized enrichment scores (NES), nominal p-values, and Benjamini–Hochberg FDR-adjusted p-values. Pathways with FDR-adjusted p-values <0.05 were deemed significantly enriched. Positive NES values indicated pathways enriched in the experimental condition, while negative NES values indicated enrichment in the control condition.

### Chromatin Immunoprecipitation (ChIP)

THP-1 macrophages were transfected with UNA-targeting or NC ASO. 24 hours post-transfection, cells were cross-linked with 1% formaldehyde (Thermo Fisher Scientific) for 10 min at room temperature and quenched with 125 mM glycine for 5 min. Cells were washed twice with ice-cold PBS. Nuclear pellets were isolated by swelling cross-linked cells in hypotonic lysis buffer (25 mM Hepes, pH 7.4, 1.5 mM MgCl2, 10 mM KCl, 0.5% NP-40, and 1 mM DTT) supplemented with 1x protease inhibitor (Sigma). Nuclear pellets were suspended in sonication buffer (50 mM Hepes, pH 7.4, 140 mM NaCl, 1 mM EDTA, 1% Triton X-100, 0.1% sodium deoxycholate, 0.5% SDS, 1 mM DTT, and 1x protease inhibitor) and incubated on ice for 10 min. Nuclear extracts were sonicated using a QSonica Q800R for 10 cycles of “30 sec ON and 30 sec OFF” at the highest voltage setting to generate 500-1000 bp chromatin fragments. Equal quantities of sheared chromatin (10 μg per IP) were diluted 1:5 in sonication buffer (no SDS) to a final volume of 1 ml and immunoprecipitated overnight at 4°C with 10 μl of RNA Pol II (Active Motif, clone 4H8, cat. 39097) antibody or isotype control IgG (Santa-Cruz SC-2025). Chromatin complexes were captured using 20 μl Dynabeads Protein G (Invitrogen) at 4°C for 3 hr. Beads were washed once with sonication buffer (containing 0.1% SDS), twice with high-salt buffer (50 mM Hepes pH 7.4, 500 mM NaCl, 1 mM EDTA, 1% Triton-X 100, 0.1% Sodium deoxycholate, 0.1% SDS), twice with LiCl buffer (20 mM Tris pH 7.4, 250 mM LiCl, 1 mM EDTA, 0.5% NP-40, 0.1% Sodium deoxycholate, 0.05% Tween-20), and once with TE buffer (10 mM Tris pH 7.4, 1 mM EDTA). Each wash was performed at room temperature for 5 min in 1 mL. Beads were captured using a DynaMag magnet (Thermo Fisher Scientific). Elution was performed by suspending beads in 100 μl of elution buffer (20 mM Tris, pH 7.4, 1% SDS, 50 mM NaHCO3, 1 mM EDTA). ChIP eluates were reverse-crosslinked at 65 °C for 4 hr, digested with Proteinase K (10 mg/ml) at 55 °C for 1 hr, and with 2 μl of RNase cocktail (Ambion) at 37 °C for 30 min. ChIP-enriched DNA was purified using PCR purification columns (Qiagen) and subjected to qPCR analysis with primers that amplify genomic regions around the transcription start sites (TSSs) of the indicated genes. Primer sequences used in ChIP studies are provided in Supplementary Table S1. All ChIP results are shown as fold enrichment in Pol II pulldown relative to isotype IgG pulldown for the respective experimental conditions.

## Data availability statement

The data that support the findings of this study are available from the corresponding author upon reasonable request.

---

**Supplementary Table S1:**
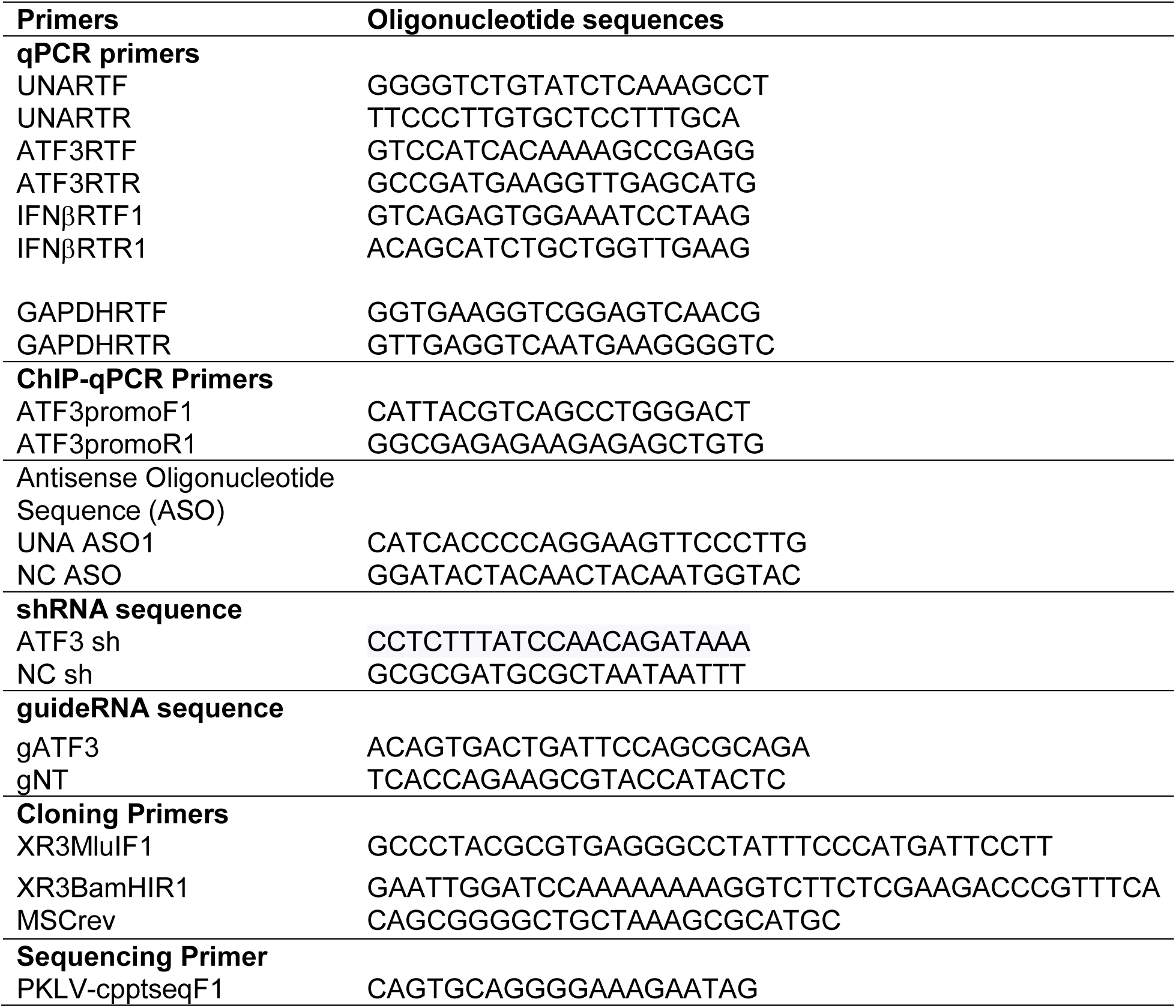
Oligonucleotide sequences.

**Supplementary Figure S1.**
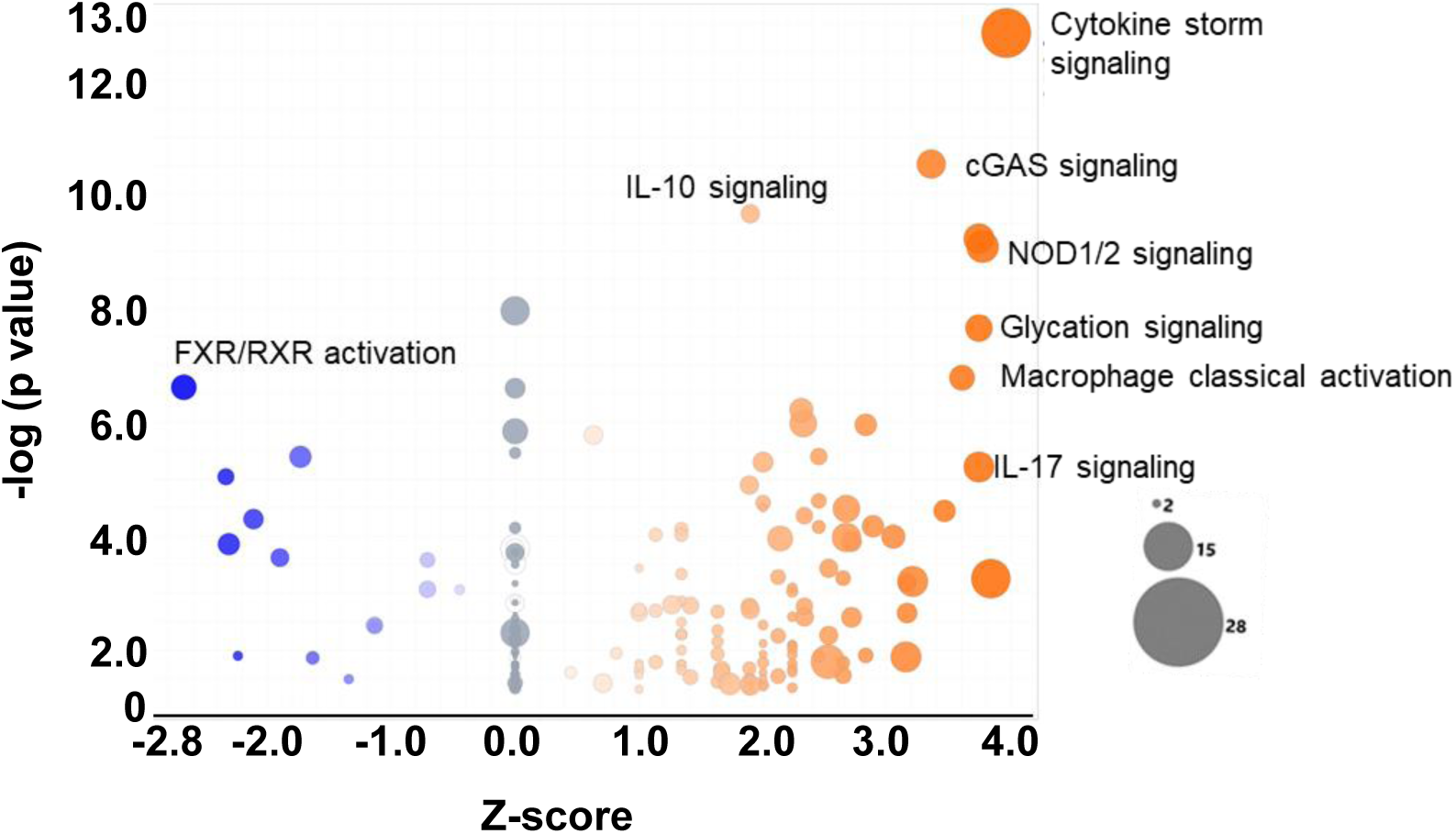
Canonical pathway analysis of EBOV-induced transcriptional responses in macrophages. Ingenuity Pathway Analysis (IPA) was performed on differentially expressed genes identified by transcriptomic profiling of EBOV-infected versus mock-infected macrophages at 24 h post-infection. The bubble plot displays significantly enriched canonical pathways, with each circle representing a pathway. Bubble size reflects the number of differentially expressed genes mapping to each pathway, and color indicates the direction and relative magnitude of pathway enrichment (blue, negative enrichment; orange, positive enrichment). Significantly regulated pathways are annotated. This analysis highlights robust activation of innate immune and inflammatory signaling programs.

**Supplementary Figure S2.**
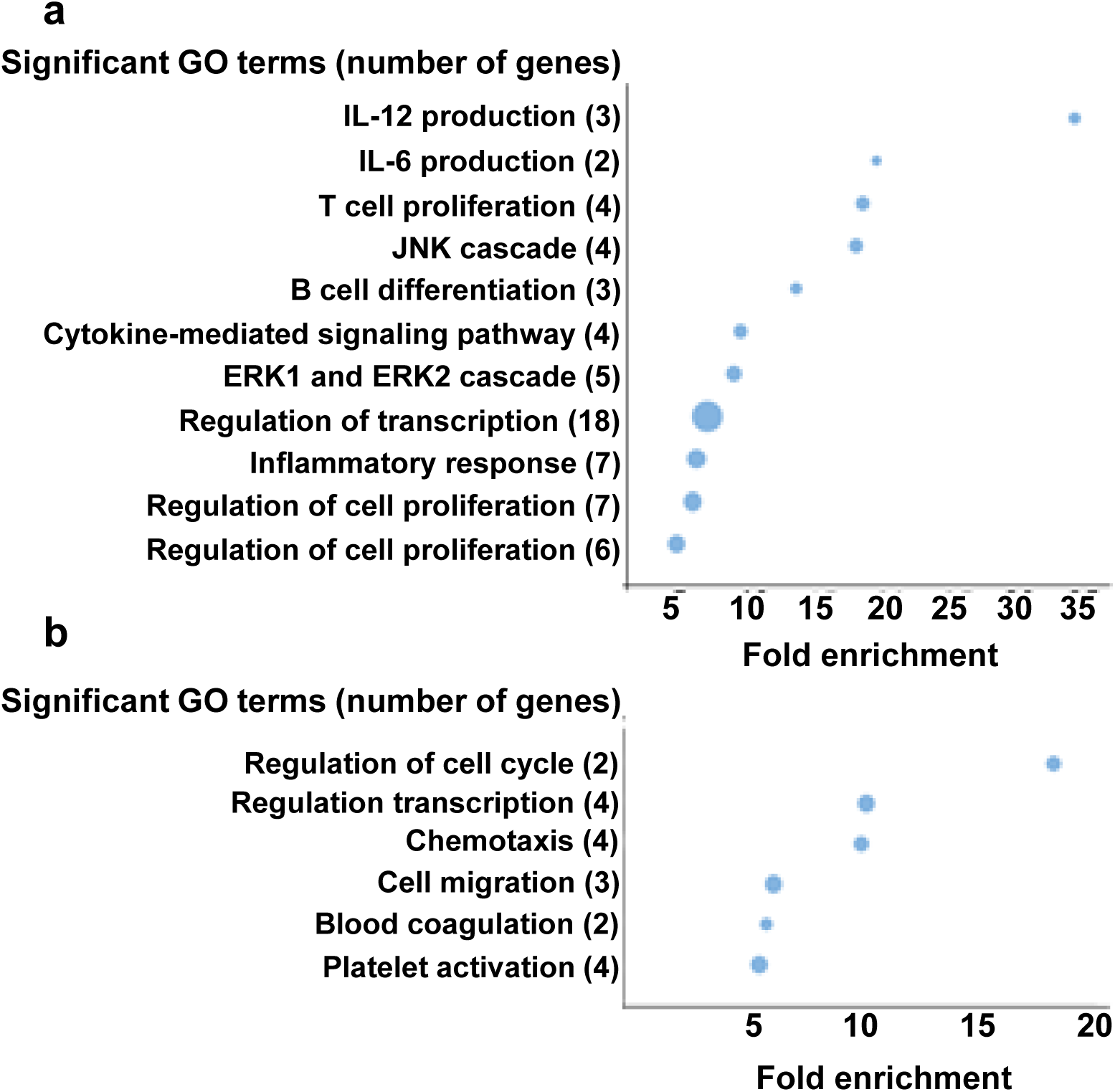
Dynamic lncRNA-mRNA Co-regulatory Networks during EBOV Infection. Protein-coding genes involved in transcriptional machinery and immune responses are co-regulated with differentially expressed lncRNAs at 24 h **(a)** and 48 h **(b)** after EBOV infection in human macrophages. Gene Ontology analysis was performed on protein-coding mRNAs co-regulated with the associated lncRNAs. Significantly enriched pathways are shown, with bubble size indicating the number of genes in each pathway, which are also indicated in parentheses.

**Supplementary Figure S3.**
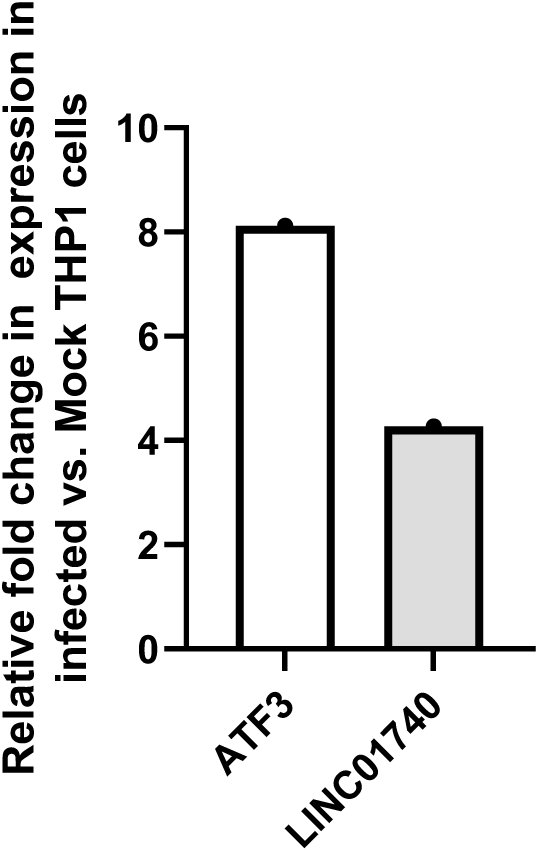
The Makona variant of EBOV induces UNA and ATF3 expression in THP-1 macrophages. RNA-seq analyses from a previously published study showed significant induction of UNA and ATF3 in PMA-differentiated THP-1 macrophages infected with the Makona variant. Expression of UNA and ATF3 was most significantly increased 72 h post-infection in THP-1 macrophages.

**Supplementary Figure S4.**
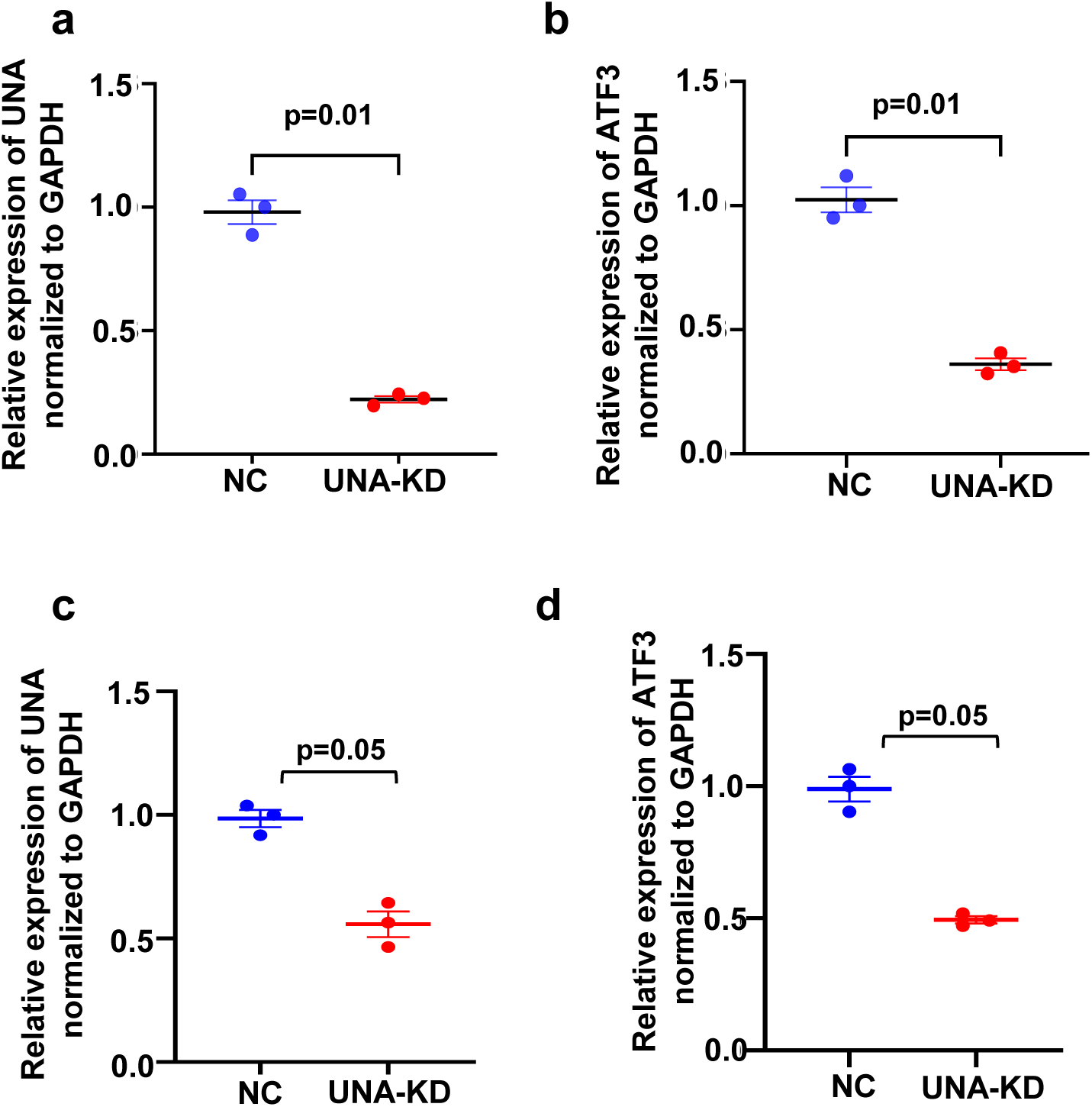
UNA knockdown reduces ATF3 mRNA expression in human cells. Cells were transfected with an antisense oligonucleotide (ASO) targeting UNA (UNA-KD) or a negative control ASO (NC). **(a)** UNA transcript levels were significantly reduced in LentiX UNA-KD cells compared with NC-transfected cells. **(b)** ATF3 mRNA was also significantly decreased in UNA-KD LentiX cells relative to NC-transfected cells. **(c, d)** Similarly, in HeLa cells, UNA-KD cells showed a significant reduction in UNA and ATF3 transcripts compared with controls (NC). Data are presented as mean ± SEM. Student’s t-test was used for statistical comparison, and a two-tailed p-value is indicated.

**Supplementary Figure S5.**
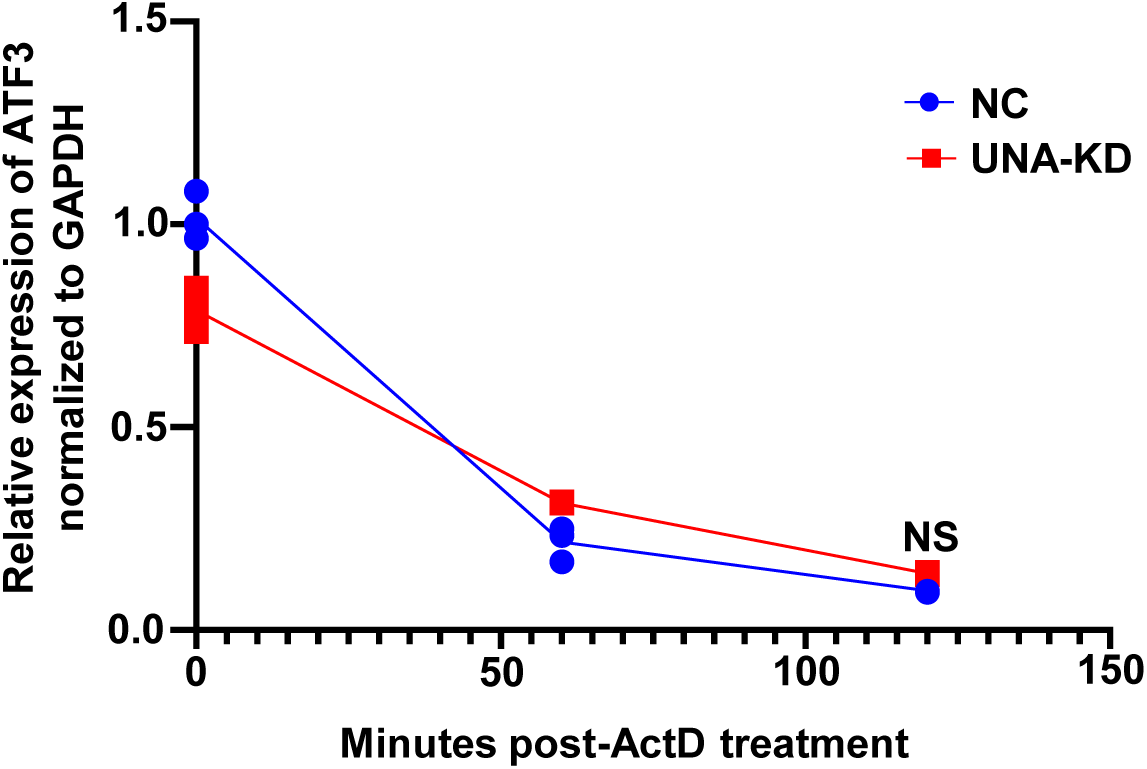
UNA knockdown did not affect ATF3 mRNA stability. PMA-differentiated THP-1 cells were transfected with an antisense oligonucleotide (ASO) targeting UNA (UNA-KD) or a negative control ASO (NC) for 24 hours. Subsequently, de novo RNA synthesis was blocked with 1 µg/mL actinomycin D. Cells were harvested at the indicated time points, total RNA was extracted, and mRNA expression was measured by qPCR. ATF3 mRNA levels at each time point were compared with those measured before actinomycin D addition. Data are presented as mean ± SEM. Student’s t-test was used for statistical comparison; NS = not significant.

**Supplementary Figure S6.**
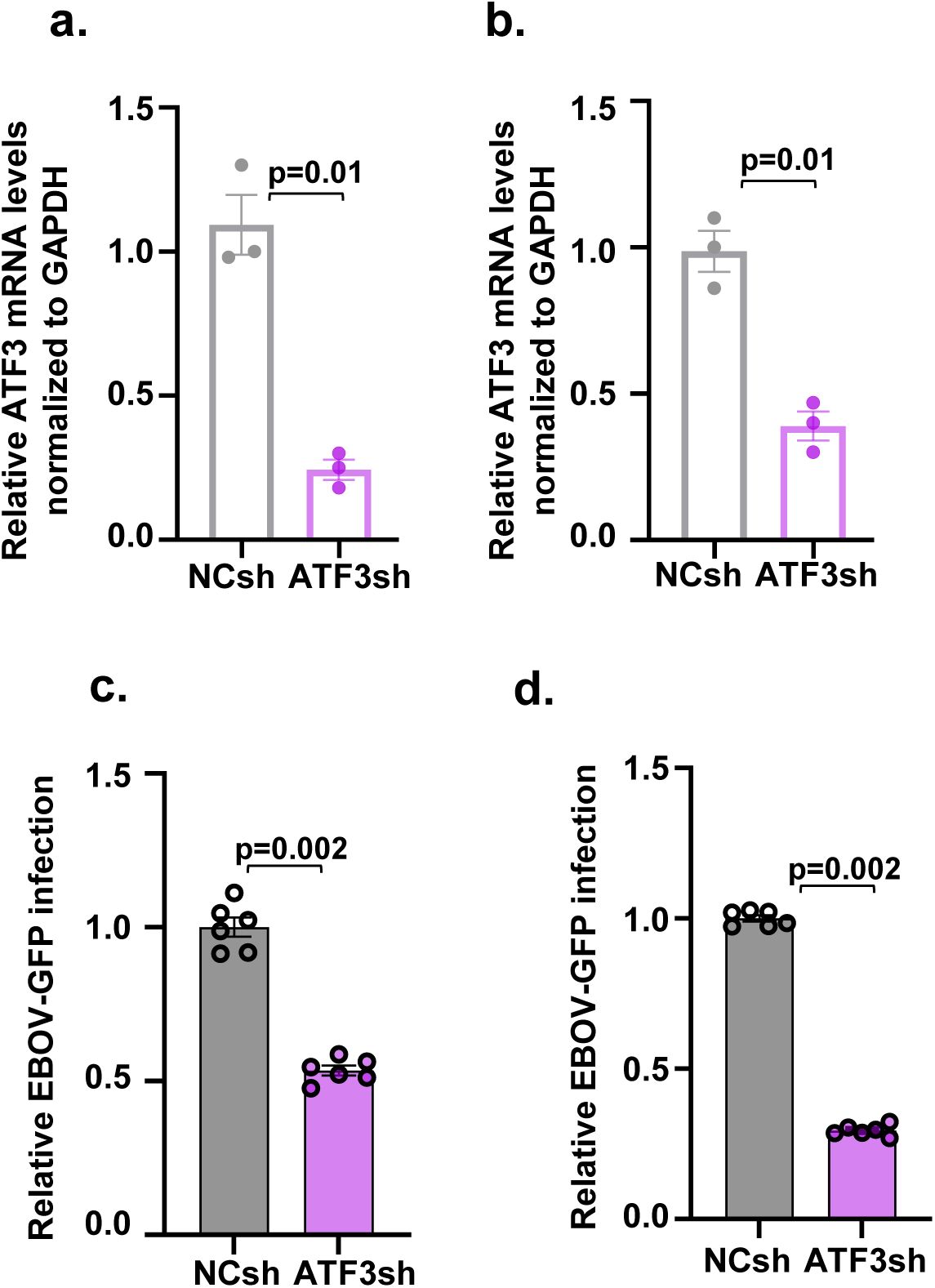
ATF3 knockdown inhibits EBOV infection in HeLa and Huh cells. **(a, b)** We used ATF3-targeting shRNAs (ATF3sh) or a negative ontrol (NCsh) to knock down ATF3 expression in HeLa (epithelial) or Huh hepatocyte) cells. We infected cells with EBOV-GFP at an MOI of 0.2 for 24 h and measured EBOV-GFP+ cells. We observed a significant reduction in EBOV-GFP+ levels in ATF3sh vs. NCsh in both HeLa **(c)** and Huh **(d)** cells. Data are presented as mean ± SEM. Student’s t-test was used for statistical comparison.

